# Carbon Fiber on Polyimide Ultra-Microelectrodes

**DOI:** 10.1101/123281

**Authors:** Winthrop F. Gillis, Charles A. Lissandrello, Jun Shen, Ben W. Pearre, Alket Mertiri, Felix Deku, Stuart Cogan, Bradley J. Holinski, Daniel J. Chew, Alice E. White, Timothy J. Gardner, Timothy M. Otchy

## Abstract

Most preparations for making neural recordings degrade over time and eventually fail due to insertion trauma and reactive tissue response. The magnitudes of these responses are thought to be related to the electrode size (specifically, the cross-sectional area) and the relative stiffness of the electrode material. Carbon fiber ultramicroelectrodes have a much smaller cross-section than traditional electrodes and thus may enable improved longevity of neural recordings in the central and peripheral nervous systems. Only two carbon fiber array designs have been described previously, each with limited channel densities due to limitations of the fabrication processes or interconnect strategies. Here, we describe a method for assembling carbon fiber electrodes on a flexible polyimide substrate that will facilitate the construction of high-density recording and stimulating arrays for acute use in peripheral nerves. Fibers were aligned using an alignment tool that was 3D-printed with sub-micron resolution using direct laser writing. Indium deposition on the carbon fibers provided a robust and reliable method of electrical connection to the polyimide traces. Spontaneous action potentials and stimulation-evoked compound responses with SNR > 10 and > 120, respectively, were recorded from a small (125 μm) peripheral nerve. We also improved the typically poor charge injection capacity of small diameter carbon fibers can be improved by electrodepositing 100 nm thick iridium oxide films, making the carbon fiber arrays suitable for electrical stimulation as well as recording.

## Introduction

With recent advances in the field of neuromodulation, it has become clear that selective chronic stimulation of the peripheral autonomic nervous system can manipulate signals passing between the central nervous system and visceral end organs to treat disease in humans (Birmingham et al., 2014; Borovikova et al., 2000; Famm et al., 2013; Koopman et al., 2016; Moore, 2015; Morris et al., 2013; Sackeim et al., 2001). Nerve interfaces typically used in this new field of bioelectronic therapy rely on centimeter-scale cuff electrode architectures to modulate activity within the cervical vagus nerve (Koopman et al., 2016; Liporace et al., 2001; Sackeim et al., 2001). Because the cervical vagus carries both afferent and efferent signals for several major organs, bulk stimulation can induce a variety of undesirable, off-target side effects (Koopman et al., 2016; Liporace et al., 2001; Sackeim et al., 2001; Verrier et al., 2016).

To provide a more precise therapeutic effect, electrodes can be positioned on target nerves closer to the end organ, where the fibers within the autonomic nerve are likely to be more specific to a single function of that organ (Bakris et al., 2012; Conde et al., 2014; Famm et al., 2013). Another approach for a precise treatment is to place the electrode closer to target fibers within the nerve with an epineurium-penetrating array. When electrodes are placed within the epineurium, electrical stimulation thresholds are lowered and effects may be localized to adjacent fascicles (Boretius et al., 2010; Tyler and Durand, 2002; Yoshida and Horch, 1993), hypothetically providing organ or function-specific modulation. Similarly, penetrating nerve interfaces are capable of making fascicle-specific or axon-specific recordings with higher spatial specificity and signal-to-noise ratio than is possible using electrodes placed on the epineurium surface (Branner et al., 2001).

Sub-cellular scale carbon fiber “ultra-microelectrodes” have been proposed to reduce insertion damage and chronic tissue response relative to conventional microelectrodes (Vitale et al., 2015; Yoshida Kozai et al., 2012). However, the challenge of building multichannel arrays from such fibers is significant as methods for micromanipulating, assembling, and making reliable electrical connections with the small-scale fibers are not yet developed. Recently, we demonstrated that multi-channel arrays can be assembled from carbon fibers using a 3D printed micro-channel funnel (Guitchounts et al., 2013). Another group has reported a linear 16 channel array containing carbon fibers mounted in grooved silicon support structures (Patel et al., 2015). The micro-patterned support structure is a key innovation that enables precise, small-scale construction. However, a significant drawback to both designs is the macroscopic scale of the carbon fiber electrode interconnect, which would make implantation within the periphery difficult.

Here, we describe a method for fabricating carbon fiber arrays with small interconnects that are appropriate for implanting in limited access areas. Though inspired by the micro-patterned carbon fiber support structures of (Patel et al., 2015), our approach adds two important innovations. The first is the use of nano-scale direct laser writing to create a micro-patterned alignment jig to support aligned carbon fibers. The second is the use of electrodeposited indium to form robust contacts between carbon fibers and copper traces embedded in polyimide thin films, allowing further miniaturization of the device relative to prior methods relying on conductive adhesives for attaching carbon fibers (Guitchounts et al., 2013; Patel et al., 2015). We tested the arrays in an acute *in vivo* preparation of a small peripheral nerve.

## Methods

### Carbon Fiber Preparation

We used 4.5 μm diameter carbon fiber threads (Grade UMS2526; Goodfellow, Coraopolis, PA) as the primary recording element of the devices. (We have previously reported the suitability of these fibers for use in recording neural activity (Guitchounts et al., 2013)). The epoxy sizing was removed from the fibers by heating them in a Paragon SC2 kiln (Paragon, Mesquite, TX) to ~400 C for 6 hrs. Fibers were then individually isolated by ‘combing’ the bundle with adhesive-backed paper (Post-It Notes; 3M Corporation, St. Paul, MN). The carbon fibers were placed in parallel on a custom-made ‘harp’ fiber holder (Figure 1a). The fiber harp was modeled using solid-modeling and design software (SolidWorks 2015; Daussalt Systèmes SolidWorks Corporation, Waltham, MA) and printed using a Form1+ 3D printer (FLGPBK02 photopolymer; Formlabs, Somerville, MA; harp design files available upon request). To prepare the harp, a strip of double-sided tape was placed on each prong of the holder and a thin strip of adhesive-backed paper was attached (adhesive side facing outwards) to the double-sided tape to allow for lifting all fibers simultaneously. Several fibers (5-10) were strung across the the harp with approximately 5 mm spacing. All of the fibers were then electrically shorted by placing conductive tape (XYZ-Axis Electrically Conductive Tape #9713; 3M Corporation, St Paul, MN) over each end of the fibers.

**Figure 1.**
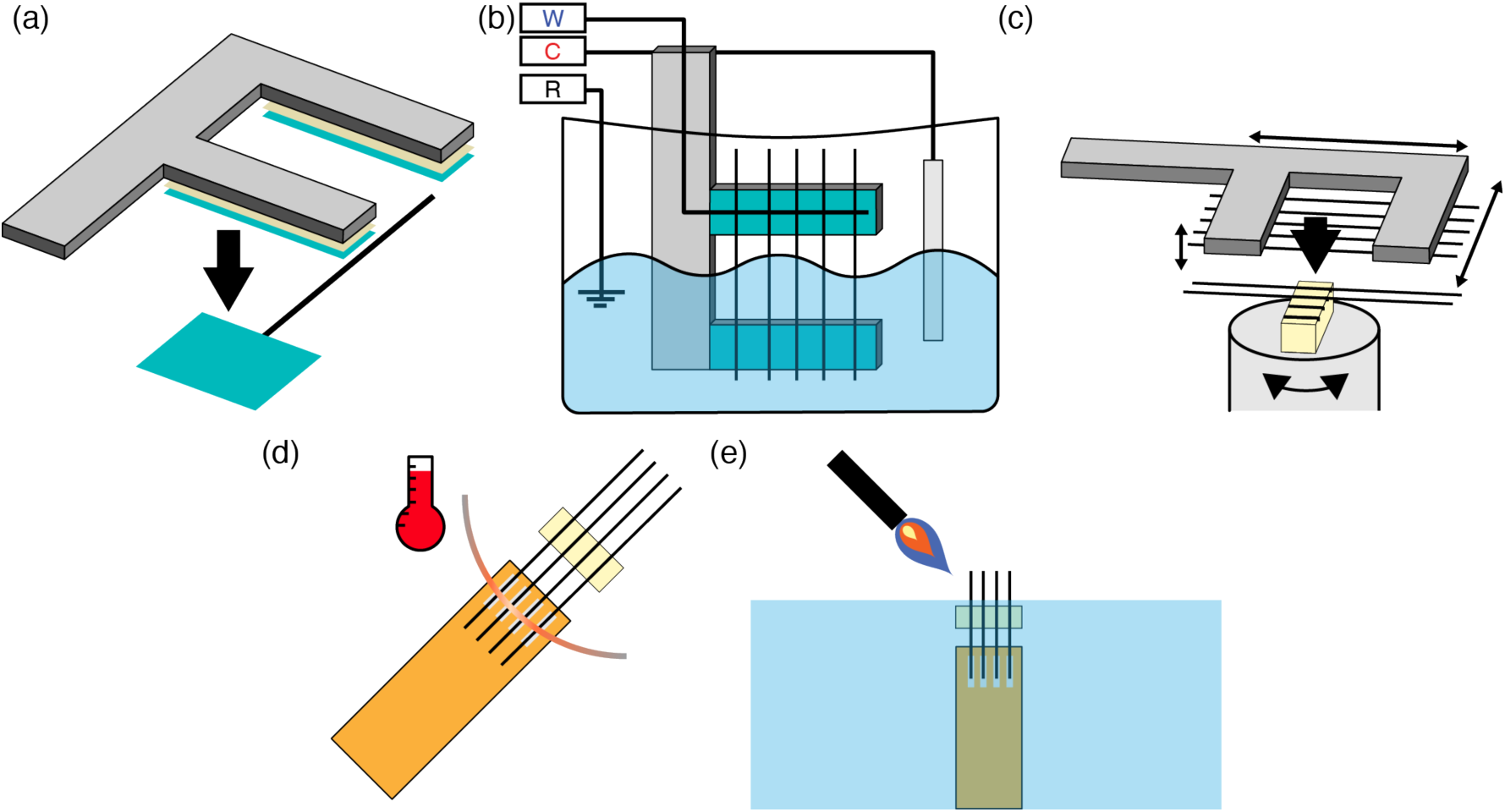
Schematic of the carbon fiber array assembly process. (**a**) First, fibers are placed on a harp-like structure to improve positioning and handling during subsequent steps. (**b**) Next, carbon fibers are partially submerged into an indium plating bath and cyclic voltage sweeps are applied to induce electrodeposition of indium onto one end of the fibers. W: carbon fiber working electrodes; C: indium counter electrode; R: Ag|AgCl reference electrode. (**c**) Using the fiber harp, the indium plated carbon fibers can be precisely positioned in a microgroove alignment clip. The alignment clip securely retains already positioned fibers, preventing accidental jostling and displacement during subsequent fiber insertions. (**d**) A polyimide lead with indium plated pads is brought in contact with the aligned, indium plated carbon fibers, and the two are electrically bonded with a fine heating element. (**e**) The carbon fibers are trimmed to length and the electrode tips of the electrodes are fire-sharpened at the air-water interface.

### Indium Electrodeposition

Using a Gamry Reference 600 potentiostat (Gamry Instruments, Warminster, PA), indium was electrodeposited onto the carbon fibers. The potentiostat was used in a 3-electrode cell configuration with Ag|AgCl as the reference electrode, thin strips of indium metal (Indium Corporation, Clinton, NY) as the counter electrode, and the prepared carbon fibers, electrically connected to the potentistat lead by the previously placed conductive tape, as the working electrodes (Figure 1b). The counter and reference electrodes and the lower half of the fiber harp were suspended in an indium sulfamate plating bath solution (Indium Corporation, Clinton, NY), and a constant current of -100 μA was applied (450 sec). This process resulted in the deposition of a 4-8 μm thick indium coating over the submerged section of the carbon fibers (Figure 2).

**Figure 2.**
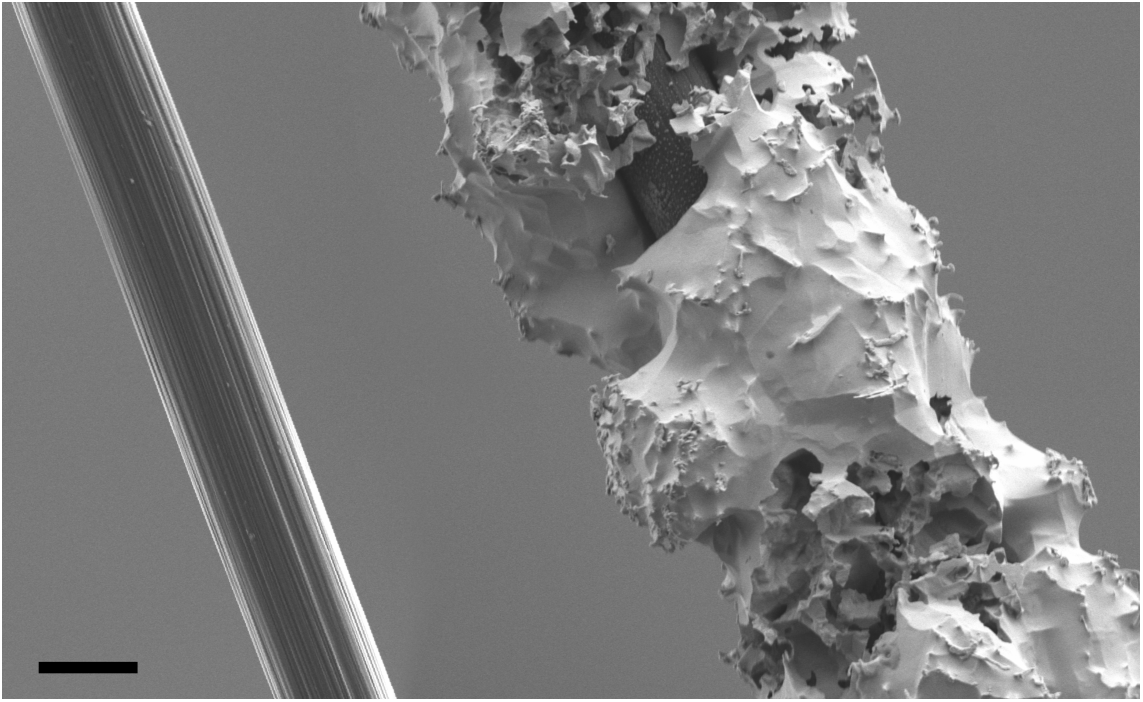
SEM micrographs of carbon fibers. (Left) Bare fiber with epoxy sizing removed. (Right) Fiber with electrodeposited indium coating; coatings were typically 4-8 μm thick. Scale bar: 4 μm.

### Polyimide Lead Design

Next, a polyimide (PI) lead (Rigiflex Technology, Inc., Anaheim, CA) was custom designed using circuit board design software (Eagle PCB; Autodesk, Mill Valley, CA). This lead, which specifies the number and spacing of individual channels in the array, served as an interconnect between the carbon fiber electrodes and an Omnetics connector (A79040 or A79042; Omnetics Corporation, Minneapolis, MN) for convenient interfacing with recording or stimulating amplifiers. The connector was soldered to one end of the lead using an SMT reflow oven (AS-5001; SMTmax, Chino, CA). The other end of the lead was prepared for carbon fiber electrode bonding by rinsing it with isopropanol and spraying off the excess with nitrogen gas. The exposed copper contacts were then placed into the indium sulfamate plating bath, and indium was electrodeposited (600 sec at -100 μA) using the plating configuration described above.

### Fiber Alignment Clip

To achieve precise, parallel alignment of the carbon fiber electrodes, we developed a 3D-printed alignment clip which allows us to place the carbon fibers, one-by-one, into microgrooves that position the fibers at a prespecified pitch (Figure 3). The alignment clip was designed using SolidWorks and fabricated from a UV-curable photopolymer (IP-DIP; Nanoscribe GmbH, Eggenstein-Leopoldshafen, Germany) using a two-photon direct laser writing system (Nanoscribe Photonic Professional GT, Nanoscribe GmbH). This system can print objects with feature sizes as small as 200 nm, allowing us to build precise structures that prevent dislodging of already placed carbon fibers while subsequent fibers are positioned in the clip. The alignment clip reported here was designed with 150 μm spacing between electrodes (matching the pitch of the PI lead), but we have also used the same methods to fabricate electrode arrays with pitch spacing as small as 20 μm (not shown). Alignment clips are incorporated into the finished electrode array (see Figures 1e, 4b), so a new clip must be fabricated for each array.

**Figure 3.**
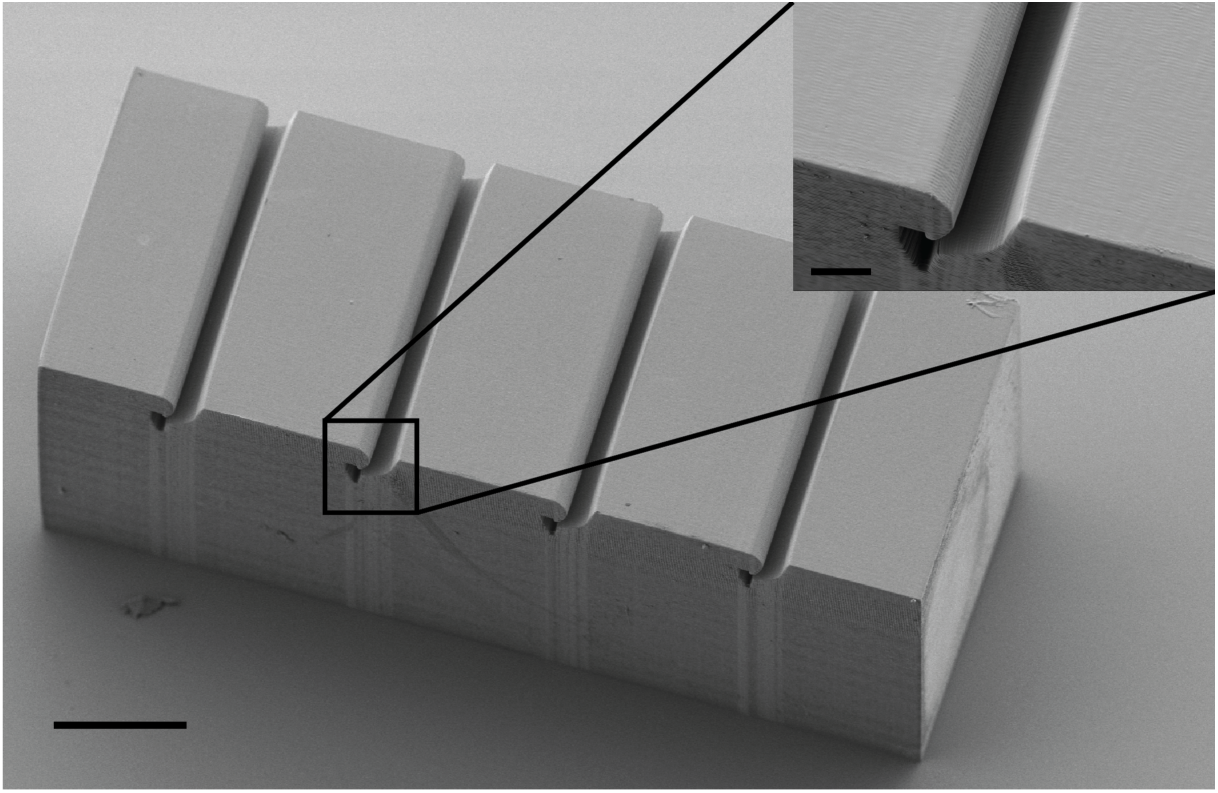
SEM micrograph of the carbon fiber alignment clip. Fibers are placed into the parallel microgrooves of this device to align them for bonding to the polyimide cable. Inset highlights the tapering of the microgroove and termination in the retaining cavity. Scale bar: 100 μm; inset scale bar: 20 μm.

### Electrode Array Assembly

The alignment clip was loosely secured to a silicon wafer or glass slide using a small amount of photo-curable cement on the (ungrooved) underside, and the wafer/slide was placed on a rotating stage (ESP301; Newport, Irving, CA) beneath a stereoscope. The fiber harp was held in a 3-axis micromanipulation stage (MP-285; Sutter Instruments, Novato, CA) that allowed for precise positioning of individual fibers, and the whole assembly was positioned above the alignment clip (Figure 1c). Carbon fibers (with deposited indium on one side) were slotted into each of the microgrooves of the alignment clip and released from the fiber harp (Figure 1c). Note that although the device demonstrated here has only four channels, the same method described could be used to construct higher channel count arrays with higher densities. A prepared polyimide lead was positioned underneath the four carbon fibers such that the indium-coated sections of the fibers were in contact with the indium-coated copper pads of the polyimide lead (Figure 4a). A 76 μm diameter platinum wire was used as a heating loop element by passing a 1 A current through it to raise its temperature above the indium melting point (~155 C). The wire was brought near the fibers and lead pad interface, melting the indium and electrically bonding the fibers to the lead (Figure 1d). Photo-curable cement was applied over the fibers and alignment clip to provide mechanical support to the assembly (Figure 4b), and the device is detached from the slide or wafer with foceps.

**Figure 4.**
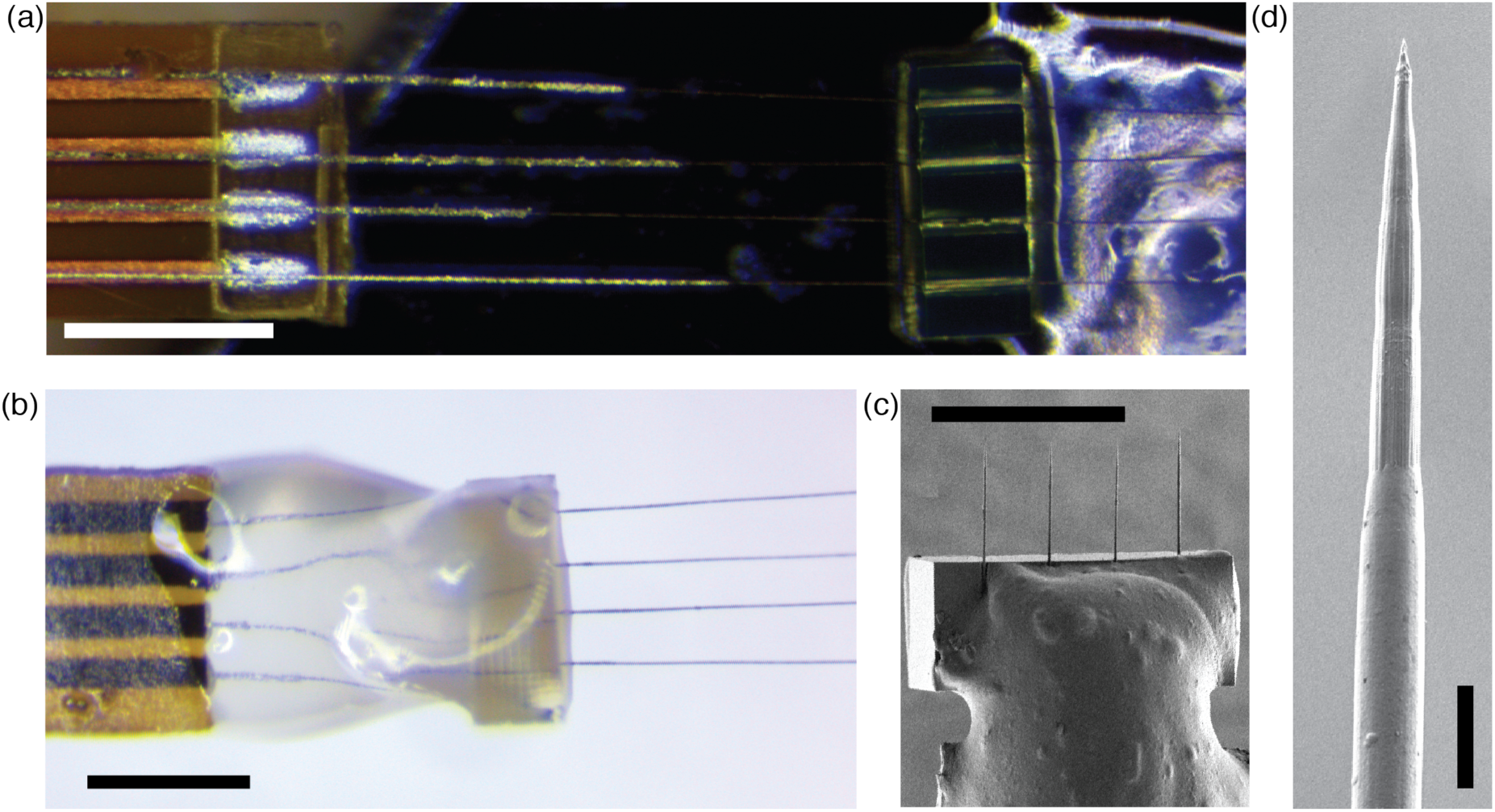
Final preparation of the carbon fiber electrode array. (**a**) Polyimide lead (left) placed underneath four aligned carbon fibers supported by an alignment clip (right). The silver-colored electrodeposited indium is visible on both the fibers and the lead pads. Scale bar: 500 μm. (**b**) Assembled array viewed from the opposite side. The alignment clip and carbon fibers were mechanically supported with photo-curable cement (clear material). Scale bar: 500 μm. (**c**) SEM micrograph of the fire-sharpened carbon fiber tips mounted in the alignment clip. Scale bar: 500 μm. (**d**) SEM micrograph of the tip of a fire-sharpened carbon fiber. Note the smooth transition to the parylene-coated section at the bottom of the image. Scale bar: 10 μm.

### Parylene Insulation and Fire-Sharpening

The contacts of the Omnetics connector were covered with masking tape, and a 1 μm layer of Parylene-C (di-chloro-di-p-xylylene; Uniglobe Kisco, White Plains, NY) was vacuum deposited over the entire device using a SCS Labcoater (PDS-2010; Specialty Coating Systems, Indianapolis, IN). The integrity of the insulation was confirmed via bubble testing of the first devices. To expose the recording tips, the fibers were fire-sharpened to the desired implantation length using a previously described air-water interface method (Guitchounts et al., 2013) (Figure 1e). This process both removed the parylene insulation and sharpened the blunt ends of the fibers to fine tips (Figure 4c-d). An Ag|AgCl reference wire was soldered to the Omnetics connector, and electrode impedances were measured in phosphate buffered saline solution (PBS, i.e. 0.9% saline) using a NanoZ impedance tester (Neuralynx).

### Iridium Oxide Electrodeposition

To reduce impedance and increase charge-injection capacity, the fire-sharpened electrodes were plated with electrodeposited iridium oxide film (EIROF) by methods previously described for gold or platinum substrates (Cogan, 2008; Meyer et al., 2001). Briefly, an electrodeposition solution was prepared by dissolving a 4 mM IrCl_4_ in 40 mM oxalic acid, followed by gradual addition of 340 mM K_2_CO_3_ to a final pH of 10.3. This solution equilibrated over a 14-day period before use in electrodeposition. A three-electrode cell comprising the fire-sharpened carbon fiber tips as working electrodes, an Ag|AgCl reference electrode, and a large surface area platinum counter electrode was used for all electrochemical characterization and deposition. Baseline cyclic voltammery (CV) to estimate the charge storage capacities (CSC) and electrochemical impedance spectroscopy (EIS) measurements were made for each electrode in PBS. For measuring CSC, cyclic voltammetry sweeps were between -0.6 V and 0.6 V at 50mV/s (Figure 5); impedance was characterized at every decade between 100 Hz and 1 MHz at 10 mV RMS. Iridium oxide films were deposited using 50 CV cycles at a 50 mV/s sweep rate between -0.05 V and 0.5 V. The electrodes were then rinsed with de-ionized water, and CV and EIS were remeasured in PBS. The time integral of the area under the CV curve was used to estimate the CSC (Cogan, 2008).

**Figure 5.**
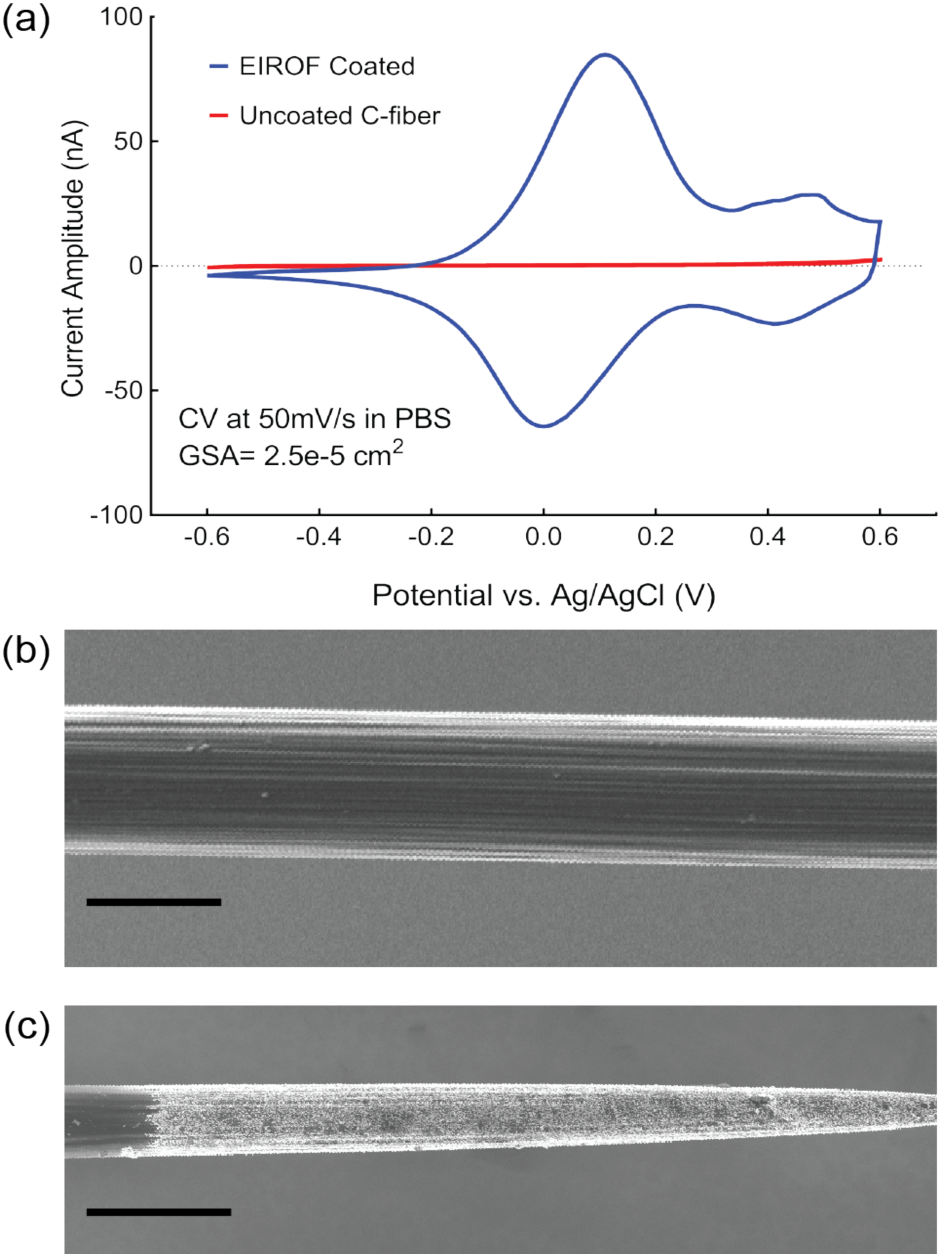
Electrodeposition of iridium oxide films (EIROF) can modify carbon fiber electrical properties. (**a**) Cyclic voltammetry curves, measured in PBS, for a carbon fiber electrode before (red) and after (blue) EIROF coating. (**b-c**) SEM micrograph of carbon fiber before (**b**) and after (**c**) fire-sharpening and EIROF coating. Note uniform iridium oxide coating over the entire electrode tip. Scale bars in (**b**) and (**c**): 4 μm and 10 μm, respectively.

### Surgical Procedure

The care and experimental manipulation of all animals were reviewed and approved by the Boston University Institutional Animal Care and Use Committee. The electrode array was tested in acute recording and stimulating preparations targeting the tracheosyringeal (TS) nerve (a branch of the hypoglossal nerve, nXII) in *n* = 6 zebra finches *(Taeniopygia guttata;* both male and female birds were used in these experiments). The TS nerve, which runs along the length of the songbird trachea and terminates at the primary vocal organ, has a diameter of approximately 125 μm and is composed of 99% myelinated fibers (Lissandrello et al., 2017). Prior to surgery, the birds were food and water restricted for 30-60 minutes to minimize the potential for aspiration during the procedure. All birds were anesthetized with 4% isoflurane dissolved in oxygen and delivered via a head mask at 0.5 L/min. Once breathing slowed and the animal was unresponsive to a slight toe pinch, the isoflurane concentration was reduced to 2% for the duration of the experiment. Meloxicam (120 μL, 1% Metacam in PBS), a nonsteriodal anti-inflammatory drug, was administered IM. The bird was placed in a supine position and a small firm clay block was placed underneath the neck for support. Feathers were removed from the lower head, neck and upper chest region to cleanly expose the neck. Betadine antiseptic solution (5% povidone-iodine) and 70% ethanol were successively applied to sterilize the exposed skin. A 15-20 mm incision in the skin of the neck was made, and the tissue blunt dissected to expose the trachea. Sutures were placed in the skin of the lateral edge of the incision and retracted from the body to better expose the implant site. Connective tissue surrounding the nerve was blunt dissected away, and two sections of the nerve (each 3-4 mm and approximately 5-8 mm apart) were isolated from the trachea. Each isolated nerve section was elevated off the trachea with a custom-made, two-prong tungsten hook. Using a stereotactic manipulator, a carbon fiber array was inserted into the more caudal segment of isolated nerve through the epineurium and into the center of the nerve (Figure 6). In some experiments, the array was held in position using only the manipulator; in other experiments, the array was stabilized in the nerve using photo-curable cement. Tissue dehydration during the procedure was minimized through generous application of PBS to the nerve and surrounding tissues. At the conclusion of the experiment, the animal was euthanized with a lethal dose of Euthasol (Virbac, Fort Worth, TX).

**Figure 6.**
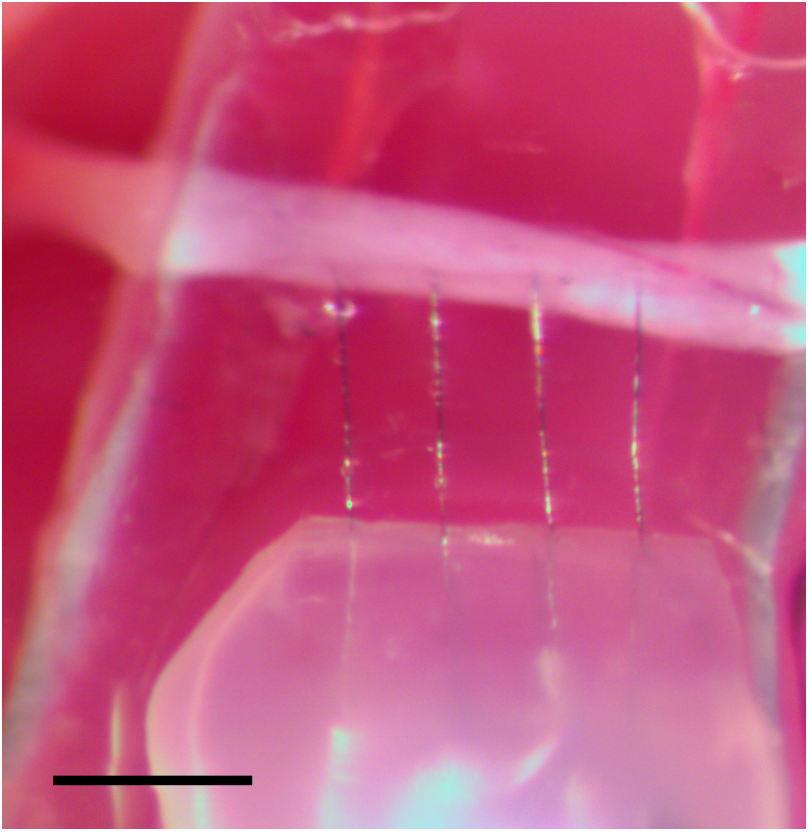
Carbon fiber array (bottom) implanted in the tracheosyringeal nerve (horizontal white band). The fire-sharpened carbon fibers easily penetrated the intact epineurium sheath. Scale bar: 300 μm.

### Electrophysiology: Recording

After nerve isolation, the array was inserted into the nerve to record electrical activity. To minimize current leakage and improve the isolation from the body, the PBS immediately surrounding the implant was displaced with a small amount of mineral oil. The Ag|AgCl reference wires were placed in contact with the tissues of the cervical cavity at some distance from the nerve. Two types of nerve activity were recorded: spontaneous, respiratory coupled activity (*n* = 2 birds) and stimulation-evoked responses (*n* = 3 birds). We tested the electrode arrays with two different neural recording amplifiers: an Intan RHD 2000 with a 16-channel unipolar input headstage (Intan Technologies, Los Angles, CA) and a Medusa Preamplifier (TDT, Tucker-Davis Technologies, Alachula, FL). The Intan system was used to record spontaneous respiratory-coupled activity, while the TDT system was used to record stimulation-evoked responses. The recorded raw voltage traces were bandpass filtered in the range 0.3 Hz-7.5 kHz (for the Intan-based recordings) or 1 Hz-10 kHz (for the TDT-based recordings). Sampling rates were 20 kSa/s and 24 kSa/s for the Intan and TDT recordings, respectively. At the conclusion of the recording session, the nerve was transected rostral to the recording array to verify that the recorded activity was from neural sources and was not instead motion-induced or caused by other recording artifacts. Recording sessions lasted approximately 30 minutes.

### Electrophysiology: Tungsten Hook Electrode Stimulation

A PlexStim stimulator (Plexon, Dallas, TX) was used to provide bipolar, current-controlled stimulation to the nerve. Custom MATLAB software (The Mathworks, Natick, MA) was written to control the Plexon stimulator (software available upon request). The two-prong hook elevating the more rostral nerve section was used as a stimulating electrode for recording evoked responses through the carbon fiber array. All stimulation was biphasic, 100 μs/phase, (100 ms interpulse intervals) at amplitudes spanning 2 μA to 805 μA.

### Electrophysiology: 4-channel array stimulation

In n=1 bird, an EIROF coated 4-channel carbon fiber array was implanted into the more rostral nerve segment and used for intrafascicular stimulation. The nerve was stimulated through two of the four channels on the electrode array (i.e., in a bipolar configuration). Again, all stimulation was biphasic, 100 μs/phase (100 ms interpulse intervals) at amplitudes spanning 38 μA to 309 μA. Evoked responses were recorded with a nanoclip recording interface (Lissandrello et al., 2017) placed 8 mm caudal to the stimulation site.

## Results

### Evaluation of Assembly Method

We used indium alloy electrodeposition on polyimide leads and carbon fibers as a method of making robust electrical connections between typically unsolderable materials. Electrodeposition of indium onto the polyimide cables could be confirmed by visual inspection under magnification for evidence of a color change from copper-brown to silver (Figure 4a). The carbon fibers were similarly visibly altered after electrodeposition, with a light silver layer visible on the electrode surface. To further characterize the electrodeposition on the fibers, several were imaged with a scanning electron microscope and showed a layer of indium alloy about 4-8 μm thick (Figure 2).

We assayed the design and performance of the alignment clip by slotting several carbon fibers into the clip, lightly brushing fibers placed in the block with a sharpened tungsten pin, and counting the number of fibers displaced. Early versions of the alignment clip had wedge-shaped or rectangular indentations, which allowed too much freedom of motion for positioned fibers and high displacement counts. After much experimentation, we converged on a design with wide channels that taper into a narrow neck and an oblong retaining cavity (Figure 3). This design allowed for easy positioning of fibers in the wide grooves but very secure retention once the fiber was moved down the tapered neck. The high resolution of the direct laser writing process enables fabrication of fine features and ultrasmooth surfaces that minimize the potential for rough clip edges obstructing fiber placement.

The strength with which the carbon fibers were bonded to the polyimide lead pads was tested by gently displacing the polyimide lead with respect to the alignment clip restrained fibers to verify that the fibers did not come free. In the assembly of six 4-channel arrays, only one fiber (i.e., ~4%) slipped from the indium bond and this was easily remedied by additional heat application. This suggests that indium soldering is a reliable method for electrically bonding materials that are generally unsuitable for other, higher-temperature soldering processes.

The fire-sharpening process that we used reliably removed the parylene insulation, exposing between 75-100 μm of the bare carbon fiber (as estimated by confocal microscopy, n = 24 fibers) and leaving uniformly sharpened tips of approximately equal length (Figure 4c). The high heat of the process has the added benefit of sealing the remaining parylene tightly against the fiber (Figure 4d), assuring smooth insertion into the tissue and minimizing the possibility of liquid penetration underneath the insulation that could lead to degredation of stimulation and recording efficacy. This process consistently produced an carbon fiber electrodes with an impedance of approximately 1 MΩ at 1 kHz (Guitchounts et al., 2013). We note that this impedance measurement included whatever (likely small) impedance due to the carbon fiber-indium-polyimide lead pad bonding.

Though impedances on the order of a megaohm are suitable for making stable neural recordings in the central (Guitchounts et al., 2013) and peripheral (Lissandrello et al., 2017) nervous systems, the voltages required to induce suprathreshold currents through such a device typically exceed the hydrolysis threshold (~0.8 V), resulting in electrode fouling and tissue damage. We found that EIROF coating the fire-sharpened tips (Figure 5c) decreased the electrode impedance 10-fold and increased the charge injection capacity from ~0.1 mCcm^-2^ (uncoated fiber) to ~25 mCcm^-2^ (~100 nm thick film; Figure 5a), consistent with previous reports (Cogan, 2008; Meyer et al., 2001).

### Acute Electrophysiology

The carbon fiber array was tested in acute recording and stimulation preparations in the TS nerve of anesthetized songbirds. Our previously reported histological analysis of the 125 μm diameter TS nerve showed that it consists of a single fascicle containing ~1000 axons, 99% of which are myelinated (Lissandrello et al., 2017).

Upon initial insertion of the array into the nerve, we observed brief, cyclic bouts of robust multiunit activity on all recording channels that appeared synchronized with the expiratory phase of the bird’s breathing (n = 2 birds; example shown in Figure 7). This seeming respiration-locked pattern persisted for the duration of the recordings and ceased only when the TS was transected rostral to the array and the bird continued to breathe, confirming that the responses were of neural origin and not a result of respiration-induced motion artifact. It has been previously suggested that the labia of the syrinx, the vocal organ innervated by the TS nerve, may partially obstruct the respiratory pathway when fully relaxed and that normal (i.e., silent, nonsinging) respiration may require periodic tensing of syringeal muscles to minimize such obstruction (Goller and Suthers, 1996; Mitra and Fee, 1998). This spontaneous, respiration-locked activity may be evidence of the neural activity necessary to induce such labial tensing.

**Figure 7.**
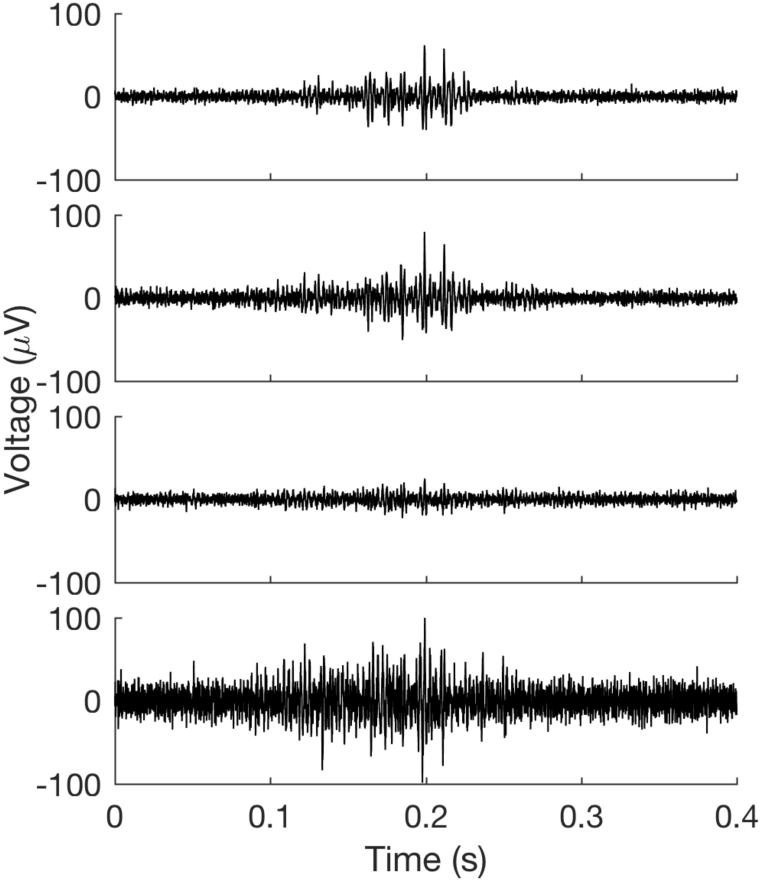
Example recording of spontaneous, respiration synchronized activity from the TS nerve. Each voltage trace shows simultaneously recorded activity from one of the four electrode array channels.

Using tungsten-hook stimulating electrodes placed rostal to the array, we recorded stimulation-evoked responses in the TS nerve in n = 3 birds (Figure 8). Repeated stimulations at a constant suprathreshold intensity reliably evoked responses of consistent shape and amplitude (Figure 8, black), indicating that the array was likely stable and recording activity of the same subpopulation of axons over the duration of the experiment. To confirm that evoked responses were of neural origin, the nerve was transected between the stimulating hooks and the recording array. We observed immediate and complete cessation of activity on all channels – even with stimulation intensity was increased to more than 50 times threshold (Figure 8, red).

**Figure 8.**
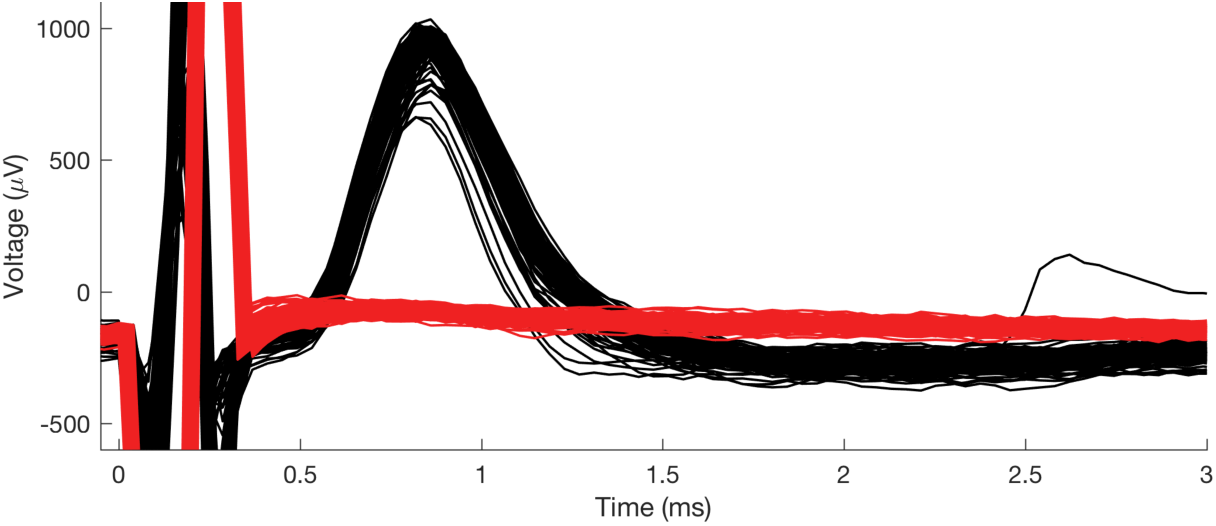
Stimulation-evoked responses recorded with the carbon fiber array. (Black) The rostral end of the TS was stimulated with bipolar tungsten hook electrodes (biphasic, 100 μs/phase, 15 μA pulses; n = 116 trials). (Red) Following transection of the TS nerve between the stimulation and recording electrodes, evoked response was abolished even at very high stimulation amplitudes (biphasic, 100 μs/phase, 805 μA pulses; n = 550 trials). The stimulation artifact is shown at *t* < 0.4 ms.

Finally, we demonstrated the efficacy of using the EIROF coated carbon fiber array for intra-nerve stimulation in n = 1 bird. With the array implanted rostral to a nanoclip recording device (Lissandrello et al., 2017), we stimulated the TS nerve with biphasic pulses using two of the carbon fibers as a bipolar electrode. We found that gradually increasing the stimulation insensity produced a graded evoked response with peaks appearing between 0.8 and 1 ms after stimulation onset (Figure 9a). Post hoc analysis showed that with increasing stimulation currents, response amplitudes sigmoidally increased until asymptoting at 150 μA and 40.1±12 μV response for currents >150 μA (Figure 9b).

**Figure 9.**
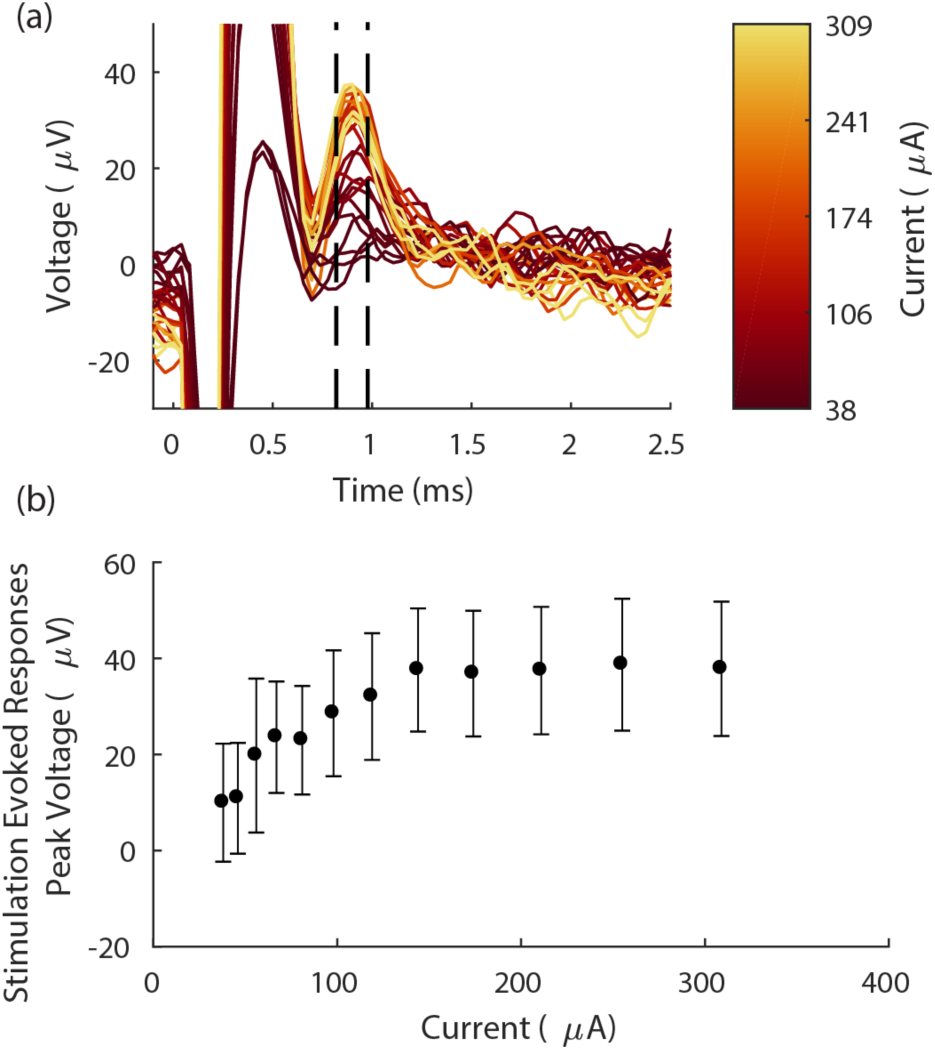
Stimulation with EIROF coated carbon fiber electrodes evokes robust, graded responses. (**a**) Carbon fiber array evoked response amplitudes increased with increasing stimulation intensity. Each trace shows the mean evoked voltage response in the nerve over n = 20 trials. Responses are aligned (at t = 0) to the onset of stimulation pulse (biphasic, 100 μs/phase); color indicates stimulation current. Dashed black lines denote window of response shown in (b). Stimulation artifact appears at 0 < t < 0.6 ms. (**b**) Mean peak amplitude of the evoked response over a range of stimulation intensities, measured between 0.8-1 ms after the stimulation onset. (Time window denoted by the dashed lines in (a)). Each data point shows n = 20 trials; error bars indicate standard deviation.

### Conclusions and Discussion

We describe a novel and flexible ultra-microelectrode array fabricated with indium metal as a key substrate for attaching fine carbon fibers to traditional metalized electrical pads. The small form factor of both the polyimide lead and carbon fiber electrodes allows for high density packing of multiple electrodes within a small implant footprint. The stiffness of the fire-sharpened carbon fibers affords easy penetration of biological tissue, including the epineurium sheath of fine peripheral nerves, offering the opportunity to make high quality intra-nerve recordings at unprecedented small scales. Furthermore, by electrodepositing a thin layer of iridium oxide on the carbon fiber tips to increase their baseline charge injection capacity, the array can also be used as a multipolar, intra-nerve stimulating electrode. We demonstrate its versatility and utility in an acute peripheral nerve preparation, but the fine geometry makes it an attractive electrode for integration with ultralight microdrives (Fee and Leonardo, 2001; Otchy and Ölveczky, 2012; Yamamoto and Wilson, 2008) to make chronic recordings in small freely-behaving animals.

With this optimized design and assembly, individual carbon fibers can be slotted into the alignment clip and firmly clasped while additional fibers are positioned. Thus, the alignment clip design is a key innovation for working with fine carbon fibers – both for the device reported here as well as for future devices that require the precise alignment of multiple fibers within a small area. In addition, the carbon fiber harp can be attached to a motorized stage to position fibers in the alignment clip, eliminating a manual assembly process that might otherwise limit how small and densely packed an array could be made.

The carbon fiber ultra-microelectrode arrays reported here differ from commercially available electrode arrays in a number of respects — the most obvious is the small scale. Though assembly on a flat polyimide lead constrains the array geometry to a planar array (i.e., either one or two rows of electrodes), the flexibility and precision of the polyimide fabrication offers the opportunity for electrode densities that far surpass the limits of other leading commercial devices (e.g., ~10-15 μm spacing vs 125 μm for the Neuronexus Buzsaki64 and 400 μm for the Blackrock Utah array). Similarly, in most commercially available arrays the electrode tips are small but is the result of a steeply tapering shank that can be 50-100 μm at the base (e.g., the Utah arrays have sharp tips but are 80 μm in diameter at the base (Jones et al., 1992)). These large electrode cross-sectional areas can result in significant tissue damage and gliosis at the site of implant (Biran et al., 2005; Polikov et al., 2005) – a problem that is compounded when multiple electrodes are placed in a small area. The carbon fibers used here have a very small diameter (<5 μm) not only at the fire-sharpened tip, but along the entire shaft of the electrode. As a result, carbon fiber microelectrodes provoke minimal reactive tissue response and incidental damage in chronic implantation preparations (Yoshida Kozai et al., 2012). In addition, carbon fiber is available with a variety of thicknesses, stiffnesses, and surface smoothness profiles enabling customization of fiber types that are best suited for implant into particular tissue types or experimental preparations.

Once the required tooling is set up and the technique is mastered, assembly time for a 4-channel array is approximately 2 hours, including all steps from indium deposition through fire-sharpening. However, there are significant savings in per-array assembly time to be had in batch producing larger numbers of arrays (e.g., fibers or polyimide leads for multiple arrays can be indium plated in a single step, multiple arrays can be parylene coated at once, etc.). Should a large number of devices or devices with a large number of electrodes be desired, then a search for suitable, scalable manufacturing processes could be undertaken.

Though we assayed the array in acute recording and stimulating preparations, the length of these experiments suggests the potential for using these multichannel carbon fiber devices in chronic studies of peripheral nerve function. One of the key remaining challenges for using this device in a chronic preparation is developing a method for securing the array on a nerve that can withstand the movement and jostling any implant in the viscera would encounter. We have recently reported a micro-scale nerve cuff for securely positioning instrumentation on fine nerves, a ‘nanoclip’ (Lissandrello et al., 2017), that is fabricated with the same direct laser writing process used to make the fiber alignment clips. Given the similarity in process and materials, it may indeed be possible to incorporate these distinct innovations into a single device fabricated in a two-stage printing process (i.e., assemble carbon fiber arrays as described here, then print the nanoclip structure around the array).

## Acknowledgements

This work was supported by an NIH grant (U01NS090454-01), GlaxoSmithKline’s Bioelectronics R&D Unit (now: Galvani Bioelectronics), the Boston University Photonics Center, and the Boston University College of Engineering. We thank Dawit Semu for expert husbandry and care of the animals involved in this experiment.

